# Split Cas12a protospacer engineering enables ultra-specific, PAM-free detection

**DOI:** 10.1101/2025.07.30.667643

**Authors:** Gabriel Lamothe, Felix Veillette, Iwe Idorenyin, Camille Bouchard, Kelly Godbout, Yaoyao Lu, Joel Rousseau, Ariel Corsano, Keith Pardee, Jacques P. Tremblay

**Affiliations:** Department of Molecular Medicine, Laval University, Quebec, Quebec, G1V 0A6, Canada; Centre de recherche du CHU de Québec, Quebec, Quebec, G1V 4G2, Canada; Department of Pharmaceutical Sciences, Leslie Dan Faculty of Pharmacy, University of Toronto, Toronto, ON, M5S 3M2, Canada; Department of Mechanical and Industrial Engineering, University of Toronto, Toronto, ON, M5S 1A1, Canada

## Abstract

CRISPR-Cas12a is a programmable, RNA-guided endonuclease that has revolutionized biotechnology, with applications in genome engineering and diagnostics. To induce nuclease activity, Cas12a must first interact with the target dsDNA duplex by associating with a short protospacer adjacent motif (PAM) in the sequence. In this study we have split this target duplex to create PAM-proximal and PAM-distal duplex regions, which has allowed us to regulate trans-cleavage activity when these regions are included in combination or separately. These observations on Cas12a activity led to hypotheses into the related functional mechanisms, which we have tested and that have highlighted DNA/protein interactions during Cas12a complex assembly that were not otherwise apparent. Selective destabilization of the nucleic acid complexes appears to drive greater reliance on the Cas12a protein for complex stability. We have exploited this to provide significant improvements in both structural selectivity and nucleotide specificity in PAM-proximal and PAM-distal duplex regions, respectively. The result is an architecture that shows promise as a PAM-free ultra-specific platform to resolve single nucleotide polymorphisms.

**GRAPHICAL ABSTRACT:** 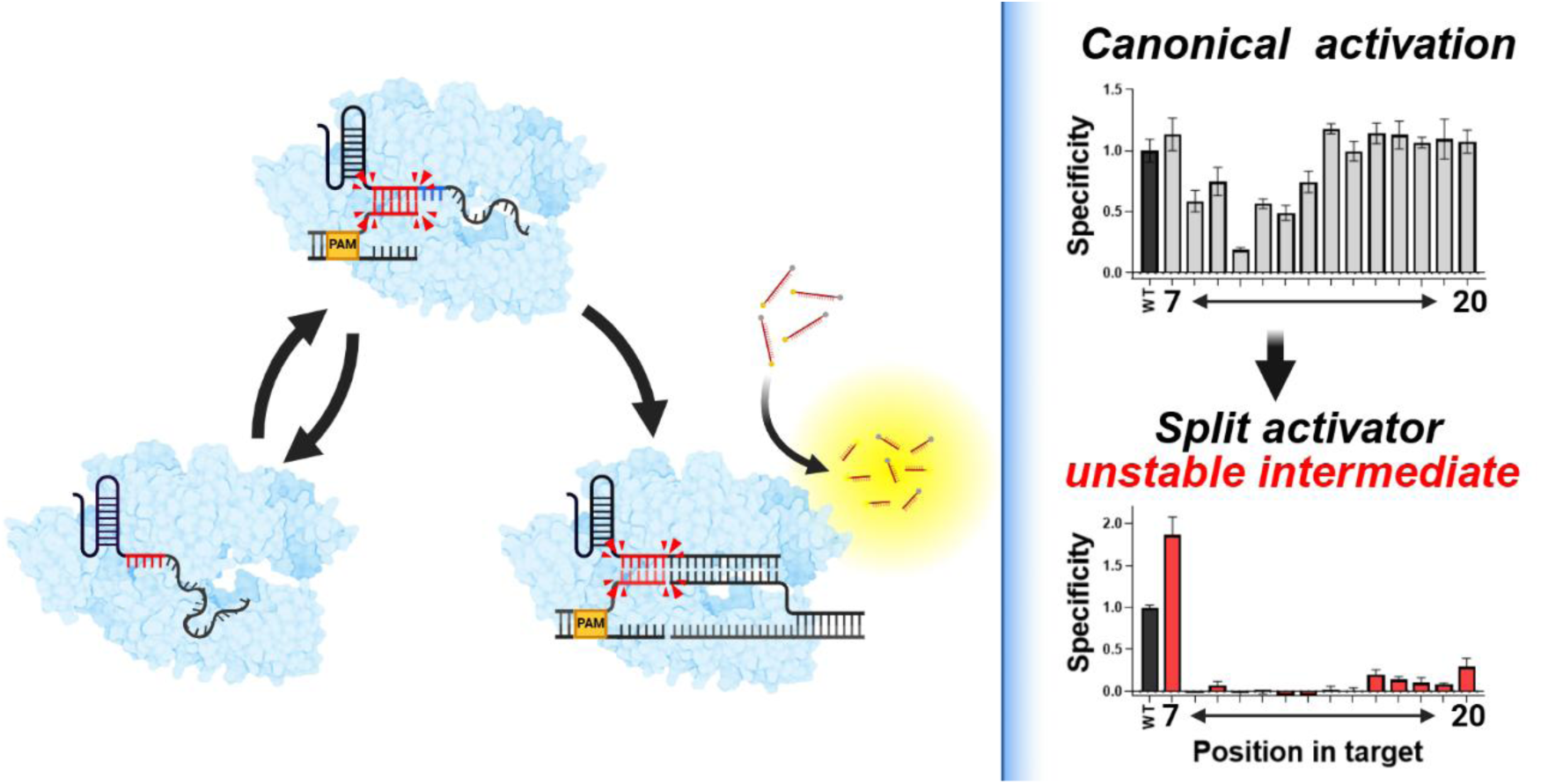

## INTRODUCTION

Clustered regularly interspaced short palindromic repeats (CRISPR)-associated protein 12 (Cas12) is primarily an RNA-guided, DNA-targeted endonuclease. Like all CRISPR proteins, it is derived from the acquired immune system of prokaryotes. It evolved to record previous infections by integrating sections of the parasitic, exogenous genetic factors (spacers) from invading bacteriophages or plasmids, into the bacteria’s genome.(1,2) These genetic factors are approximately 20 nucleotides in length and are known as protospacers.

In the case of Cas12a, these protospacers are adjacent to T-rich protospacer-adjacent motifs (PAMs) which are recognized by an RNA-guided Cas12a and used to unwind the double stranded target DNA.(3,4) Given that the PAM (5’-TTTV-3’) is required to unwind the first bases of a double-stranded activator sequence, Cas12a does not require a PAM when its activator is single-stranded.(3,5) Once the first bases are unwound, Cas12a begins hybridizing with the first ∼5 bases in its crRNA’s programmable section which are structured in an open and solvent-exposed conformation in a process known as R-loop initiation (Supplementary Figure S1A, S1B).(6)

Successful annealing leads to R-loop elongation, a process in which the crRNA displaces the non-target strand (NTS) in the double-stranded target DNA and anneals to the target strand (TS) (Supplementary Figure S1C). Once the 20^th^ base in the crRNA’s programmable region anneals to the TS, the Cas12a caps the formation of the R-loop by inserting a large aromatic residue, effectively signaling that the Cas12a has reached its activation state (Supplementary Figure S1D).(6,7) Once in this state, the nuclease site in the RuvC domain is exposed, allowing it to cleave the NTS, which is followed by cleavage of the TS by the same nuclease site, in a process referred to as cis-cleavage(8). This is followed by the release of the PAM-distal section of the DNA. At this time, Cas12a remains bound to the PAM-proximal DNA, allowing the protein to remain in its active state wherein its nuclease site is exposed to the solvent. This allows suspended ssDNA to enter the nuclease site, resulting in their indiscriminate cleavage in a process known as trans-cleavage.

Given the ease with which the crRNA can be reprogrammed to target new genes, Cas12a has been used with great success in gene editing(9–11), genetic screening(12), gene circuits (13), transcription regulation(14,15), intracellular imaging(16), and diagnostics(5,17). These unique properties have been leveraged to create a new category of low-burden nucleic acid amplification tests(5,18,19) which have significant advantages, but that have had some long standing challenges. First, as Cas12a is a primarily a DNA-targeting enzyme, it does not recognize RNA, a common substrate and important biomarker in many pathologies(20–22). In addition, dsDNA targets require a PAM (5’-TTTV-3’) sequence, which significant limits the utility of the system(5). Finally, the enzyme exhibits low fidelity for single-nucleotide polymorphisms, with researchers often having to combine natural and artificial mutations to improve the on-target resolution of their system.(5,23)

To expand the scope of Cas12a’s capabilities, recent work in the field has focused on altering the structure of the activator sequence. By introducing a break into the TS, we are left with the PAM-proximal (Pp) and the PAM-distal (Pd) activator sequences (Figure 1A).(20,24–26) These “split” activator-based systems have transformed CRISPR/Cas12a into an effector for DNA nanotechnologies and bioengineering, allowing the protein to engage in logic operations(25,27,28), constitutional dynamic networks(29), time-controlled photo-activation(29), and the programmable release of cargo(25). In support of Cas12a’s newfound relevance in the field of diagnostics(5,23), split activators have also enabled Cas12a to detect proteins, enzymes(25,28,29), and ions(30), as well as expanded its range of nucleic acid substrates to include PAM-less RNA(20,26,31). However, despite these remarkable advances, the specificity of split activator systems for single nucleotide polymorphisms is generally only slightly improved compared to the wild-type CRISPR/Cas12a system.(20,27,28) Furthermore, the majority of studies to date have focused on targeting single-stranded DNA and RNA,(20,25,27,28,30,31) with relatively little attention placed on double-stranded DNA,(24,29) despite the importance of this format in applications.

**Figure 1:**
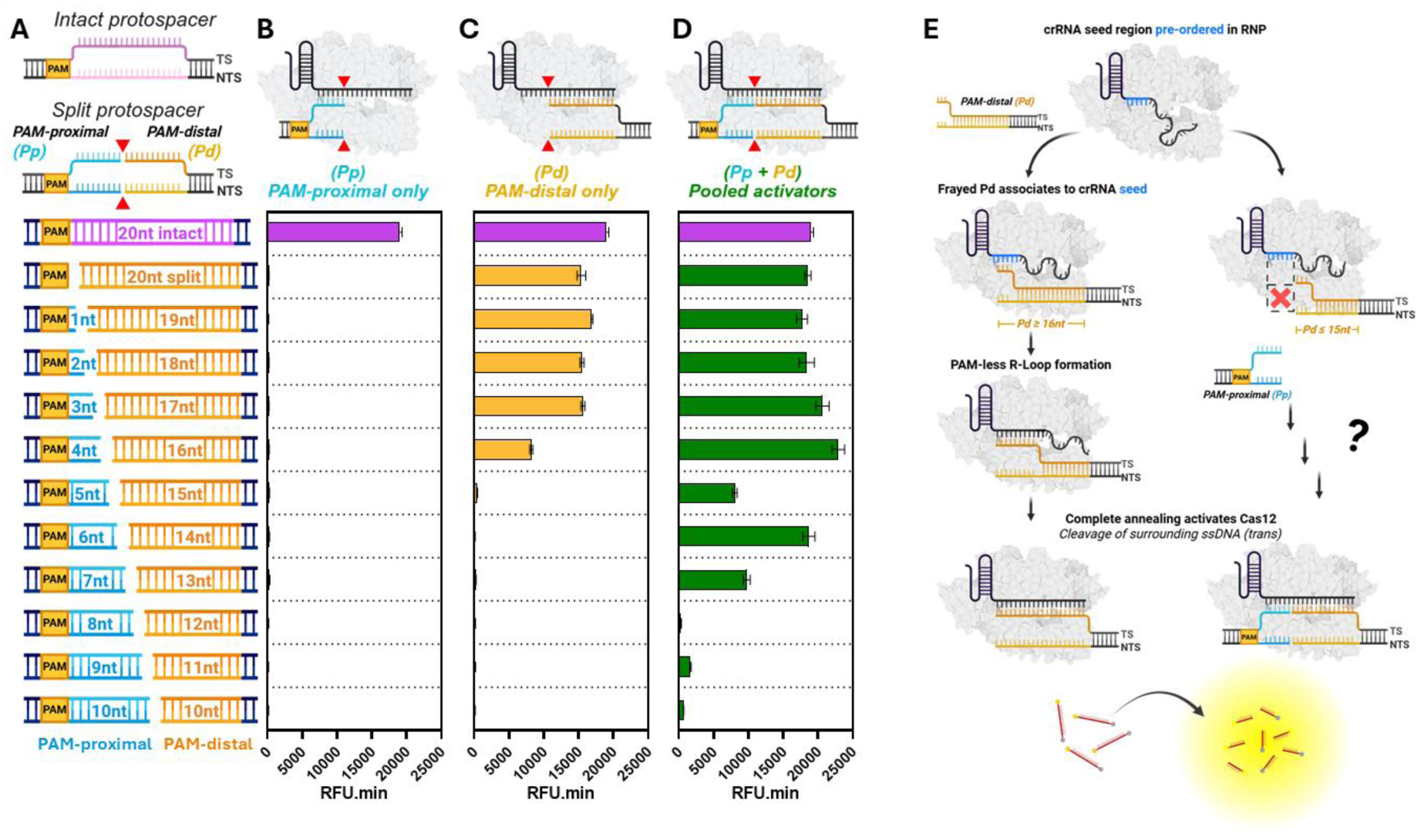
The position of the double-stranded break modulates Cas12a activity. **(A)** Cas12a activators are composed of the target (20nt) protospacer which anneals to the crRNA and the protospacer adjacent motif (PAM) which is required to unwind the duplex. Introducing a double-stranded break in the activator yields PAM-proximal and PAM-distal sequences. Breaks were introduced in the ten bases adjacent to the PAM. Trans-cleavage activity of Cas12a was measured over time when providing **(B)** PAM-proximal (Pp) duplexes only, **(C)** PAM-distal (Pd) duplexes only, and **(D)** both the Pp and Pd in combination. Conditions were benchmarked against an intact activator. Background-subtracted fluorescence intensity (n=3) was measured over 60 minutes at intervals of 3.6 minutes. Values represent total area under the curve (RFU.min) ± Standard error. **(E)** Cas12a can associate to PAM-less protospacers (Pd=20nt) provided at least the last of the first five bases adjacent to the PAM are present (Pd > 15 nt). These five bases correspond to the “seed” region in the crRNA which are pre-ordered and solvent-exposed to initiate hybridization to the target strand.(6) Graphics created in https://BioRender.com.

It is through this lens that we set out to better understand the assembly of split Cas12a systems. We began by first analyzing the response of Cas12a to split duplexes of different lengths, allowing us to focus our next steps on design that provides strong activity in the presence of both duplexes and no activity in the absence of either. We then built on a previously proposed assembly model(24) by including new conditions, integrating our findings into the original model and showing key interactions that had not been previously elucidated. Furthermore, while previous work with split activators suggested that chimeric DNA/RNA guides (crXNA) could not be used in split Cas12a systems,(20) we leveraged our improved split activator length optimizations, and structural evidence for the interaction between the guide and the Cas12a,(32) to rationally design new a crXNA-guided system. The resulting design was then leveraged to study the function of the PAM in mediating the assembly of the intermediate, ternary complex. Finally, using this selectively destabilized ternary complex, we demonstrate an improved base resolution of Cas12a for single-nucleotide polymorphisms, achieving ultra-specific signalling in almost every position of the PAM-less DNA strand. Combined, we anticipate that our optimization for split duplex lengths and the proposed mechanism for assembly of split activators will help to guide future developments in this field, with implications for diagnostics, logic operations, and DNA nanotechnologies.

## MATERIAL AND METHODS

### Reagents

EnGen^®^ Lba Cas12a (Cpf1) was purchased from New England Biolabs (M0653T, NEB, USA). NEBuffer^TM^ r2.1 was purchased from New England Biolabs (B6002S, NEB, USA). RNase inhibitor, Murine was purchased from New England Biolabs (M0314L, NEB, USA). DNAseALERT^TM^ Substrate was purchased from Integrated DNA Technologies (11-04-02-04, IDT, Coralville, IO, USA). Nuclease-Free Duplex Buffer was purchased from Integrated DNA Technologies (11-05-01-12, IDT, Coralville, IO, USA).

### Nucleic acid design and preparation

ssDNA and Guide RNAs (crRNA and crXNA) were synthesized by Integrated DNA technologies (IDT, Coralville, IO, USA). ssDNA from IDT was resuspended in IDT Nuclease-free Duplex Buffer. PAM-proximal (Pp) sequences were annealed with the target strand (TS) and the non-target strand (NTS) at a molar ratio of 1:1 unless otherwise indicated (50 µM TS, 50 µM NTS). PAM-distal (target sequences) were annealed with TS and NTS at a molar ratio of 1:1.2 (40 µM TS, 48 µM NTS). To anneal them, samples were brought to 98ᵒC for 3 minutes, then cooled to 20ᵒC over 20 minutes and kept at 4ᵒC until used. Annealed oligonucleotides were diluted in IDT Nuclease-free Duplex Buffer with a TS concentration of 1 µM and 125 nM for the Pp and Pd respectively.

### *In vitro* trans-cleavage assays

Unless otherwise indicated, trans-cleavage assays were performed by incubating 33 µL of a solution composed of 1X NEBuffer r2.1, 1 U/µL Murine RNAse inhibitor, 75 nM LbaCas12a, 75 nM Guide RNA (crRNA or crXNA), and 45 nM Pp at 37 ᵒC for 30 minutes. Then 17 µL of a second solution composed of 1X NEBuffer r2.1, 1 U/µL Murine RNAse inhibitor, 375 nM DNAse ALERT (IDT), and 30 nM Pd dsDNA was added to the first for 50 µL total. The final master mix was composed of 1X NEBuffer r2.1, 1 U/µL Murine RNAse inhibitor, 125 nM DNAse ALERT, 30 nM Pp, 50 nM LbaCas12a, 50 nM guide RNA, and 10 nM Pd. 15 µL of this master mix was aliquoted at 4 ᵒC to three wells of a 384-well opaque, black plate for three technical replicates. Plates were sealed with a sticky lid, centrifuged at 2000g for 1 minute at 4ᵒC, and read on a TECAN infinite M1000 with λ_ex_: 536 nm and λ_em_: 556nm. Plates were incubated at 37ᵒC in the TECAN and read over the course of 90-120 minutes.

### FRET Assays

The viability of some of the Fluorescence Resonance Energy Transfer (FRET) assays were first predicted by simulating the complexes we anticipated forming (Supplementary Figure S2). Simulations were run on the AlphaFold3 server (https://alphafoldserver.com/) and included sequences listed in the Supplementary Information. Structures were deemed viable if the distance between the DNA/RNA extremities of all simulations were less than 10 nm (100 Å). Fluorescent oligonucleotides were synthesized by IDT by adding Fluorescein (FAM) on their 5’ extremities. Quenching oligonucleotides were prepared by adding their IDT’s proprietary Iowa Black quencher to 3’ extremities.

Unless otherwise indicated, FRET experiments were performed by preparing a solution comprised of 1X NEBuffer r2.1, 1 U/µL Murine RNAse inhibitor, 100 nM LbaCas12a, 100 nM synthetic crRNA, and various concentrations of annealed Pp and Pd oligonucleotides. Reaction mixtures were prepared at 68 µL and three aliquots of 20 µL were added to an untreated black with clear bottom 384-well plate. Plates were incubated at 37 ᵒC in the TECAN M1000 Pro and read with an excitation wavelength of 495 nm and an emission wavelength of 520 nm.

### EMSA Binding Assay

The EMSA binding assay was performed by preparing a solution composed of 1X NEBuffer r2.1, 1 U/µL Murine RNAse inhibitor, 500 nM LbCas12a, 500 nM synthetic crRNA, and 200 nM each of Pp and/or Pd. Reactions were incubated at 37 ᵒC for one hour before placing them at 4ᵒC. 1/6 volume of EMSA loading buffer (1X NEBuffer r2.1, 50% glycerol) was added to the samples. Samples were run on 5% PAGE gel prepared with 0.5X TBE and 5 mM MgCl_2_ with a running buffer composed of 0.5X TBE and 5 mM MgCl_2_. Samples were run at 4 ᵒC in the dark.

## RESULTS

### Splitting a double-stranded protospacer

We begin by investigating how split duplexes impact the activation of *Lachnospiraceae bacterium ND2006* Cas12a (herein referred to as Cas12a). Here, a double-stranded break was introduced immediately following the PAM and extended base by base through the first 10 bases of the protospacer (Figure 1A) and separated into two parts: the PAM-proximal (Pp) and PAM-distal (Pd). When combined, the Pp (left) and Pd (right) pairs encoded the complete 20 nt protospacer. As expected, Pp activators alone failed to initiate trans-cleavage by the Cas12a, as their hybridizing sequence was less than the critical minimum of 15 nucleotides required to activate Cas12a (Figure 1B).(5)

Next, we evaluated whether the PAM, which is used by Cas12a to melt the first bases in the protospacer, was dispensable when targeting truncated DNA duplexes. Here, a PAM-free protospacer complementary to the full crRNA spacer (Pd=20nt) was placed on the extremity of a double-stranded activator sequence (Figure 1C, S3B). Under these conditions, the PAM-less Pd activator rapidly activated Cas12a, as measured by the complete trans-cleavage of the reporter. Following recent literature, we hypothesize that DNA fraying, the transient opening of DNA termini, provided access to short ssDNA segments complementary to the crRNA (Supplementary Figure S1E).(24,33) This process would be roughly analogous to the uncoupling of the activator’s first base pairs by Cas12a in the presence of the PAM (Supplementary Figure S1B).(4) Normally, this uncoupling facilitates the hybridization between the crRNA’s first ∼5 nt, which are pre-ordered and solvent exposed(6) (Supplementary Figure S1A), and the melted ssDNA immediately adjacent to the PAM (Supplementary Figure S1B, S1E). By monitoring trans-cleavage, we found that at least one of these five bases needs to be present in the Pd duplex (Pd > 15nt) for the crRNA to associate to the rest of the duplex, supporting the hypothesis (Figure 1C,E).

Having established that Cas12a could tolerate serial truncations in its activator (Pd), up to the loss of the seed region, we next investigated the effects of introducing double-stranded breaks in the activator. Here, a Pp and a Pd, which collectively encoded both the PAM and the full 20 nt protospacer, were exposed to a crRNA-guided Cas12a, otherwise known as a ribonucleoprotein (RNP). Interestingly, Cas12a’s trans-cleavage activity, which was diminished when exposed to a truncated DNA duplex (Pd=16nt) (yellow bars, Figure 1C), could be improved when the missing section of the activator sequence was provided in the form of a Pp (4nt) (green bars) (Figure 1D, S3F). This trend was even more interesting for Pd=15nt-13nt, where there is no trans-cleavage activity without both Pp and Pd component (green bars) (Figure 1D, S3G-I). This effect was not universal and could not be leveraged to significantly restore Cas12a’s activity at all positions (Supplementary Figure S3J,K,L). Importantly, the finding that Pp and Pd are both required to induce Cas12a’s activity leaves open the question as to how the truncated activators associate to the Cas12a.

### Assembly of the split protospacer

To experimentally determine the order in which the split activators assemble with Cas12a, as well as the requirements for these associations, we tracked the formation of each complex via Fluorescence Resonance Energy Transfer (FRET) assays. In short, we labeled the DNA duplexes and the crRNA with a Fluorescein (F) and a Quencher (Q), then measured for changes in the sample’s fluorescence. These changes in FRET allowed us to monitor assembly or disassociation of the nucleic acids and Cas12a, depending on the nature of the experiment. To represent the assembly of double-stranded split activator systems, the DNA break was introduced immediately following the 6^th^ base of the protospacer. When combined, the Pp/Pd split at 6bp/14bp completely restored activity in Cas12a but led to no notable activity in the absence of either (Figure 1B-D, S3H).

Since blunt-end DNA can undergo end-to-end stacking in certain conditions, we first tested whether the split protospacer fragments could assemble independent of crRNA and Cas12a.(34,35) To test this, the Pp and Pd duplexes were labelled with FAM and the quencher respectively, so that, upon interaction, a reduction in fluorescence would be observed (Figure 2A). We found that the baseline fluorescence of the Pp-FAM (Pp-F) was not quenched in the presence of a Pd-Quencher (Pd-Q) indicating that they did not undergo spontaneous end-to-end stacking (Figure 2B). Further, neither the crRNA with a sequence complementary to the Pp and Pd, nor the Cas12a which can unwind DNA duplexes, could assemble the duplexes individually. Rather, Pp:Pd assembly required all four components (Pp, Pd, crRNA, Cas12a), as can be seen with a reduction in fluorescence (Figure 2B).

**Figure 2:**
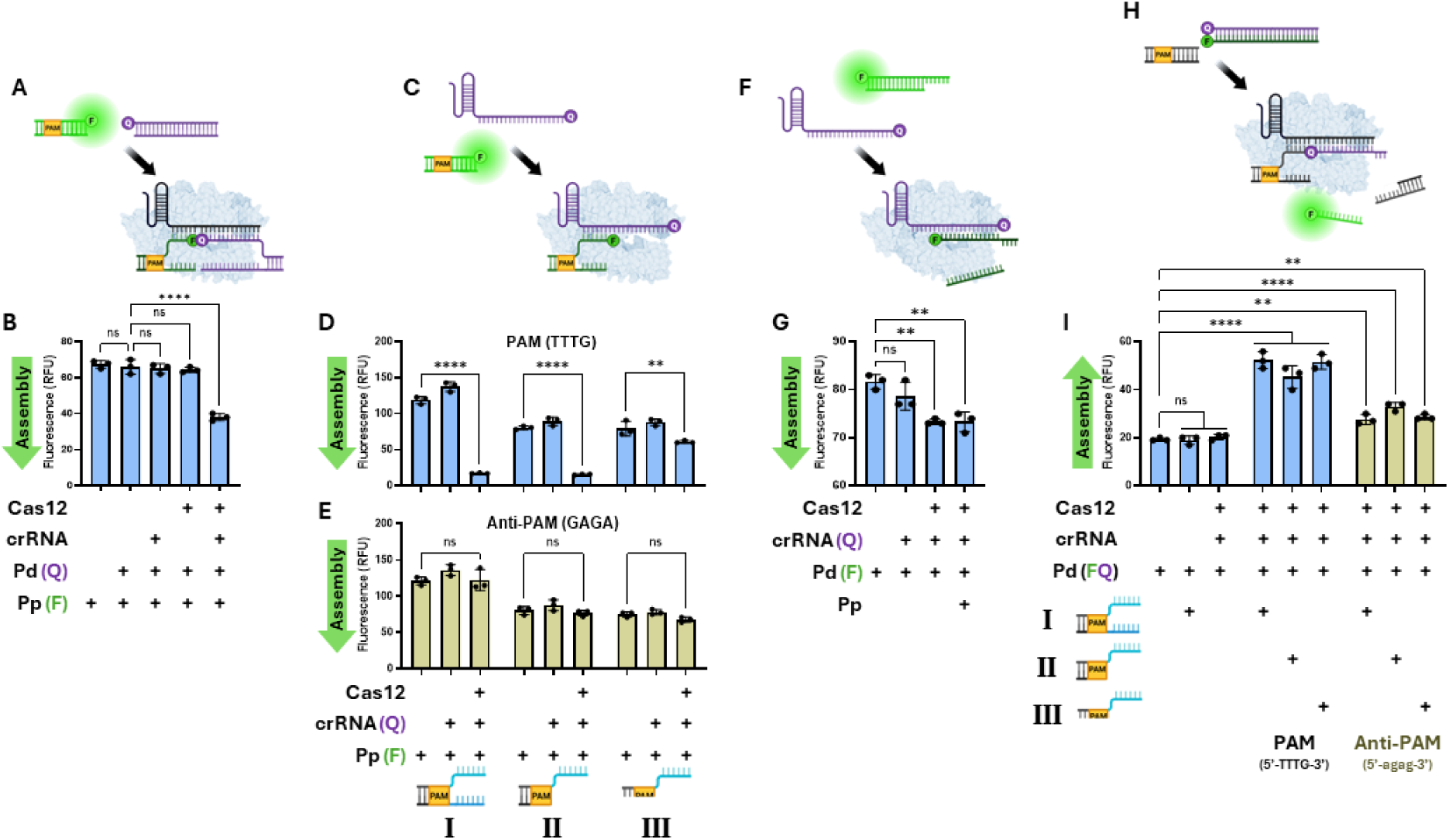
Assembly of split protospacer duplexes. To model the association of the RNP (Cas12a+crRNA) to the Pp and Pd when both are required to achieve trans-cleavage, the Pp/Pd split was placed at 6nt/14nt. **(A,B)** Assembly of Pp-F and Pd-Q (10 nM Pp-F, and 10 nM Pd-Q). (**C)** The Pp can associate to the RNP, but not the crRNA alone. The strength of association is based on **(D)** the structure and **(E)** the sequence of the PAM (20 nM Pp-F). Three structural configurations were compared, with the Pp being either I) fully double-stranded; II) double-stranded PAM, single-stranded TS; III) fully single-stranded. **(F,G)** The Pd can associate weakly to the RNP (200 nM Cas12a, 200 nM crRNA-Q, 200 nM Pp, and 10 nM Pd-F). **(H,I)** Cleavage of the Pd requires the Pp and is influenced by the sequence of the PAM (30 nM Pp, 30 nM Pd-FQ). All samples (n=3) were incubated over 30 minutes at 37 ᵒC. Values represent mean ± SD. Analyses were performed using a one-way ANOVA with a multiple comparison test where ns = not significant with *p* > 0.05 and asterisks (\**p* ≤ 0.05, \*\**p* ≤ 0.01, \*\*\**p* ≤ 0.001, and \*\*\*\**p* ≤ 0.0001) denote significant differences. Graphics created in https://BioRender.com.

Next, we sought to determine which, if either, of the split activators (Pp or Pd) would preferentially associate with the Cas12a during the formation of the active complex. Since Cas12a first interacts directly with the PAM to melt the first bases of a canonical target sequence (Supplementary Figure S1B), we reasoned that the Pp:RNP complex should be favored. To track the formation of this complex, here, we labelled the Pp with a fluorophore, and the crRNA with a quencher (Figure 2C, reduced signal indicates assembly). In the absence of Cas12a, the crRNA-Q did not quench the Pp-F regardless of whether the Pp was single-stranded, indicating that the two nucleic acid structures did not hybridize independently (Figure 2D). In the presence of Cas12a, the Pp was quenched, indicating that the protein was required to associate the two (Pp + crRNA). Interestingly, signal reduction by the Cas12a was influenced by the structure and sequence of the PAM. When the PAM was double-stranded (configurations I and II), the quenching effect was very strong (Figure 2D). In contrast, when the PAM was single-stranded (configuration III), the quenching effect was less significant. Since this difference was independent of the NTS in the crRNA’s hybridizing region (dark blue, bottom strand), this implies that the PAM may be preferentially involved in associating the Cas12a to the target DNA duplex.

To further explore the role of the PAM in mediating the assembly between the crRNA-guided Cas12a and the Pp, we replaced the PAM (5’-TTTG-3’) with an Anti-PAM (5’-GAGA-3’) (Figure 2E, reduced signal indicates assembly). Canonically, the PAM is involved in unwinding the first bases of the protospacer, facilitating the crRNA’s hybridization (Supplementary Figure S1B).(4) When the PAM is mutated, the protospacer is not accessible and the crRNA cannot hybridize. It was therefore unsurprising to find that the fully double-stranded Pp-F with an Anti-PAM did not associate to the crRNA-Q (Configuration I, Figure 2E). It was, however, surprising to find that when the hybridizing region was single stranded, the Anti-PAM still prevented the association between the Pp-F and the crRNA-Q (Figure 2E, Configurations II and III). Taken together, the effect of the PAM’s structure and sequence in the context of a split activator seems to imply that the direct DNA/protein interaction plays a significant role in associating the Pp and the Cas12a. Furthermore, this interaction is distinct from the DNA/RNA interaction between the target duplex and the guide RNA.

Recent crystallographic evidence (PDB 8Y04) from a study detailing precursor crRNA processing elucidates the structure of the RNP complexed to a double-stranded Pp with 6nt annealing to the crRNA (Supplementary Figure S4).(36) This structural evidence supports a theory that associating to the Pp improves the Cas12a’s ability to target the Pd.(24) Indeed, we demonstrated that Cas12a can target PAM-less Pd sequences, provided the pre-ordered seed (first ∼5nt) in the crRNA can hybridize (Figures 1C, S3B-G). In this crystal structure (PDB 8Y04), the crRNA bases immediately downstream of the Pp are similarly pre-ordered (bases 6-8) (Figure S5). This would support the observed activation of Cas12a along Pd targets that would otherwise not anneal to the pre-ordered bases (first ∼5nt) in the crRNA. These complimentary pieces of evidence suggest that the Pp-bound intermediate complex pre-orders the bases in the crRNA, thereby facilitating Pd-target searching.

Having established that the Pp:RNP complex can assemble independently of the Pd, and that this assembly is dictated by the PAM’s structure and sequence, we next explored whether the RNP could assemble with the Pd alone. To this end, a fluorophore and quencher were installed on the TS of the Pd and the crRNA respectively (Figure 2F). In this context, fluorescence quenching was indicative of assembly. We found that RNP-Q and Pd-F could weakly associate (Figure 2G, S6B, S6C), despite the lack of significant trans-cleavage under these conditions (Figure 1C, S3H). To confirm these results, an EMSA binding assay demonstrated that the RNP did indeed bind to the Pd-F without the Pp (Supplementary Figure S6A).(24) It was notable however that this association was weak and/or transient, as evidenced by the low level quenching during FRET experiments (Figure 2H, S6B, S6C) and co-localization during migration (Supplementary Figure S6A).

Having determined that the crRNA-guided Cas12a associates more strongly to the PAM-containing Pp than the Pd, and that this association is affected by the structure and sequence of the PAM, we next sought to complete the survey of interactions by investigating the cis-cleavage of the Pd activator. Here, we placed the fluorophore on the non-target strand and the quencher on the target strand, in this context, separation of the quencher from the fluorophore indicates cleavage and release of the fluorescent TS (Figure 2H, 2I). First, it was noted that the cis-cleavage of the Pd-FQ did not occur in the presence of only the crRNA and Cas12a, which supports the trans-cleavage findings (Figures 1C, S3H, yellow). As we can see, once a Pp was present, the Pd-FQ was cis-cleaved by Cas12a (Figure 2I), similarly supporting the trans-cleavage data (Figures 1D, S3H, green). Interestingly, the structure of the PAM did not significantly affect the cleavage of the Pd-FQ (Figure 2I, configurations I-III), despite the double-stranded PAM improving Pp-binding (Figure 2D, configurations I and II) relative to the single-stranded PAM (Figure 2D, configuration III). Instead, we see that only the sequence of the PAM significantly impacted cis-cleavage of the Pd-FQ, with the Anti-PAM reducing separation of the Pd-FQ strands (Figure 2I, yellow bars). This evidence suggests that the association between the Pp and the crRNA-guided Cas12a must be significantly impaired for Cas12a activation to cease on Pd targets.

To summarize the finds so far, these experiments add to previous work with split activators(24) in a few important ways. I)The first is that the position of the split is a key factor in whether the Pd duplex can activate the Cas12a alone (Figure 1C). II) We also provide experimental evidence that the PAM structure (or duplex configuration) and sequence dictates the assembly of the Pp:RNP complex independent of effects on Pd unwinding (Figure 2D, 2E). III) Interestingly, we also found that the RNP can associate with a short Pd independent of the Pp (Figure 2G), but that this association does not result in significant cleavage of the Pd (Figure 2I). IV) Further, we found that the Pp enables cis-cleavage of the Pd (Figure 2I) and trans-cleavage of the reporter (Figure 1D). V) Finally, when we combine these findings with previously published structural data (PDB: 8Y04), we find converging evidence that association to the Pp may pre-order the crRNA bases immediately following DNA:RNA binding, an event which likely primes the Pp:RNP for target (Pd) searching (Figure S5).

### Altering the stability of the complexes

Since the Pp:RNP association is favored in split activators (Figures S5, 3A), we hypothesized that improving the stability of this intermediate may improve its ability to target Pd, increasing activity of Cas12a. With this in mind, we first increased the GC content between the Pp and the crRNA (Figure 3B, S7); however, we did not observe improved Cas12a activity, even when the hybridizing sequence in the Pp was single-stranded to reduce interference by the NTS.

**Figure 3:**
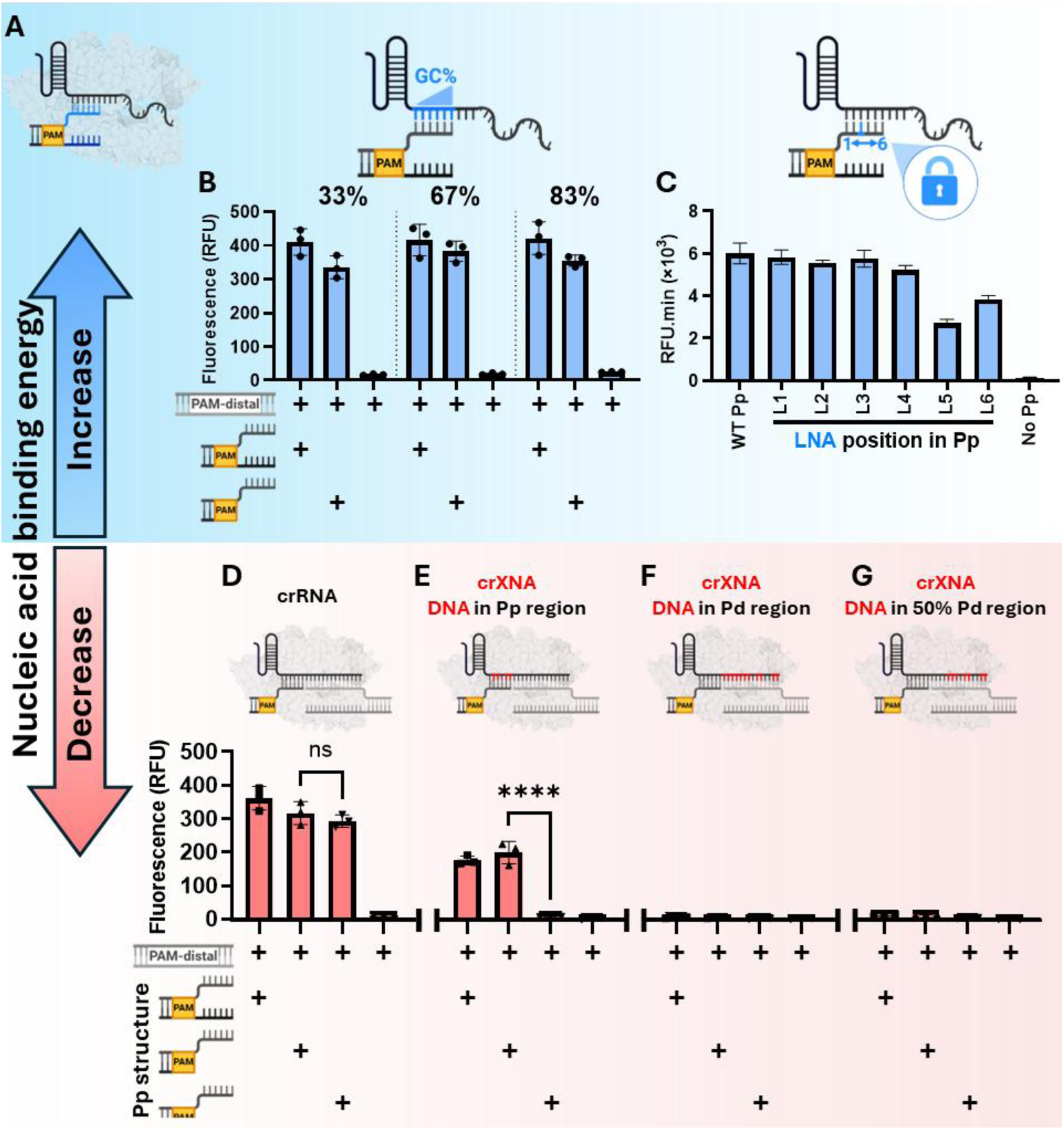
Split activator-mediated activation of Cas12a can be modulated by altering duplex stability. **(A)** Since Cas12a interacts most stably with the Pp, the crRNA:Pp duplex stability was first altered. **(B)** Trans-cleavage activity of Cas12a was measured over time using PAM-proximal hybridizing sequences with a GC content of 33%, 66%, and 83%. Conditions were benchmarked against an intact activator. Values represent mean fluorescence intensity (n=3) ± SD after incubating at 37 ᵒC over 33 minutes. **(C)** Locked nucleic acids (LNAs) were substituted into the Pp duplex’s target strand one at a time to determine Cas12a’s tolerance to the analogs. LNA-modified PAM-proximal sequences were benchmarked against non-modified (WT) Pp sequences. Background-subtracted fluorescence intensity (n=3) was measured over 30 minutes at intervals of 3.5 minutes. Finally, the trans-cleavage activity of Cas12a was measured when substituting DNA bases into the crRNA to create crXNA. Cas12a was guided by **(D)** a canonical crRNA, crXNA with DNA bases (DNA bases in red) **(E)** in the Pp-annealing region (positions 2, 3, 5 and 6), DNA bases in the Pd annealing region **(F)** (positions 7-13, 15, 16, 19 and 20) and **(G)** (positions 11-13, 15, 16, 19 and 20). Cas12a was activated using various Pp structures. Values represent mean background-subtracted fluorescence intensity (n=3) ± SD after incubation at 37 ᵒC over 100 minutes. Analyses were performed using a one-way ANOVA with a multiple comparison test where ns = not significant with *p* > 0.05 and asterisks (\*\*\*\**p* ≤ 0.0001) denote significant differences.. Graphics created in https://BioRender.com

We next introduced Locked Nucleic Acids (LNAs) into the Pp to increase the stability of the crRNA:Pp duplex.(37,38) Since LNAs have yet to be reported in split activators, we first mapped the Cas12a’s tolerance to the nucleotide analogs (Figures 3C, S8). Measurement of trans-cleavage over time revealed that LNAs in positions five and six had adverse effects on Cas12a’s activity (Figures 3C, S8H, S8I). The presence (NTS:TS duplex) or absence (single-stranded TS) of a competing NTS did not significantly impact Cas12a’s activity (Supplementary S8C-I). Furthermore, while single substitutions in positions 1-4 were tolerated (Figures 3C; S8C-G), introducing multiple LNAs at once reduced Cas12a’s trans-cleavage activity (Supplementary Figure S9). This reduction was most evident when the multi-locked Pp was fully double-stranded, likely because it was more difficult to displace the NTS and access the TS (Supplementary Figure S9C, S9D). Pre-incubating the complex for longer eliminated the difference in kinetics between the duplexed and overhanging Pp, though neither were as favorable as the unlocked Pp (Supplementary Figure S9E, S9F). Furthermore, while Cas12a is roughly as sensitive for trace quantities Pd as compared to an intact activator, locking the Pp (positions 2-4) negatively impacted the protein’s sensitivity (Supplementary Figure S10A-C,E). Given that locked nucleotides improve the stability of duplexes by shifting the DNA backbone, we conclude that Cas12a can tolerate individual but not consecutive LNAs.

Next, we wondered if destabilizing the complexes could make Cas12a more specific to alterations in the split activator sequences. Following recent literature, we decreased the binding energy of the Pp:crRNA duplex by introducing DNA bases in the guide to create chimeric DNA/RNA guides (crXNA).(39,40) To minimize the rejection of the DNA bases by Cas12a, we rationally designed the guides to avoid placing deoxyribose sugars where a previous Cas12a structural analysis (41) indicates direct contact with LbCas12a’s residues (Supplementary Figure S11). To determine the effects of weakening the hybridization between the guide and activator sequences, we compared the activity of an unmodified crRNA to the crXNAs. When guided by a crRNA, the structure of the PAM did not significantly impact the trans-cleavage activity of Cas12a (Figure 3D, S12, S13A), despite the PAM’s importance in forming the Pp:RNP intermediate (Figure 2D), which is in agreement with the cis-cleavage findings (Figure 2I). In contrast, when the Pp-annealing region of the crRNA was partially replaced with DNA, Cas12a was only significantly activated when the PAM was double-stranded (Figures 3E, S13B). This implies that weakening the nucleic acid binding energy between the Pp and the crXNA may have emphasized the PAM-Cas12a interaction (Figures 2D, 2E), making the Cas12a selective for the structure of the PAM. However, it remains unclear exactly how this was achieved. Finally, DNA substitutions in the Pd-annealing region reduced Cas12a activity, making those chimeric guides non-viable (Figures 3F, 3G, S13C, S13D).

### Improved substrate selectivity with chimeric guide

To determine if the crXNA alone made Cas12a more reliant on the PAM, we incubated the crXNA-guided Cas12a with an intact activator sequence (full 20nt), either with or without the PAM (Figure 4A). Under these conditions, just like with the crRNA (Figure 1C, S3B), the crXNA was equally active regardless of the PAM. From this, we can infer that despite the weaker binding energy, the crXNA does not inherently require a PAM to associate to its target DNA and activate Cas12a. This implies that the selectivity for the structure of the PAM in the Pp sequence (Figure 3E, 4B) may be an emergent property of a split activator. In light of these findings, we hypothesized that altering the crXNA backbone destabilized the Pp-hybridized intermediate, ultimately requiring the PAM to maintain the Pp-Cas12a association (Figure S14D, configuration III → II).

**Figure 4:**
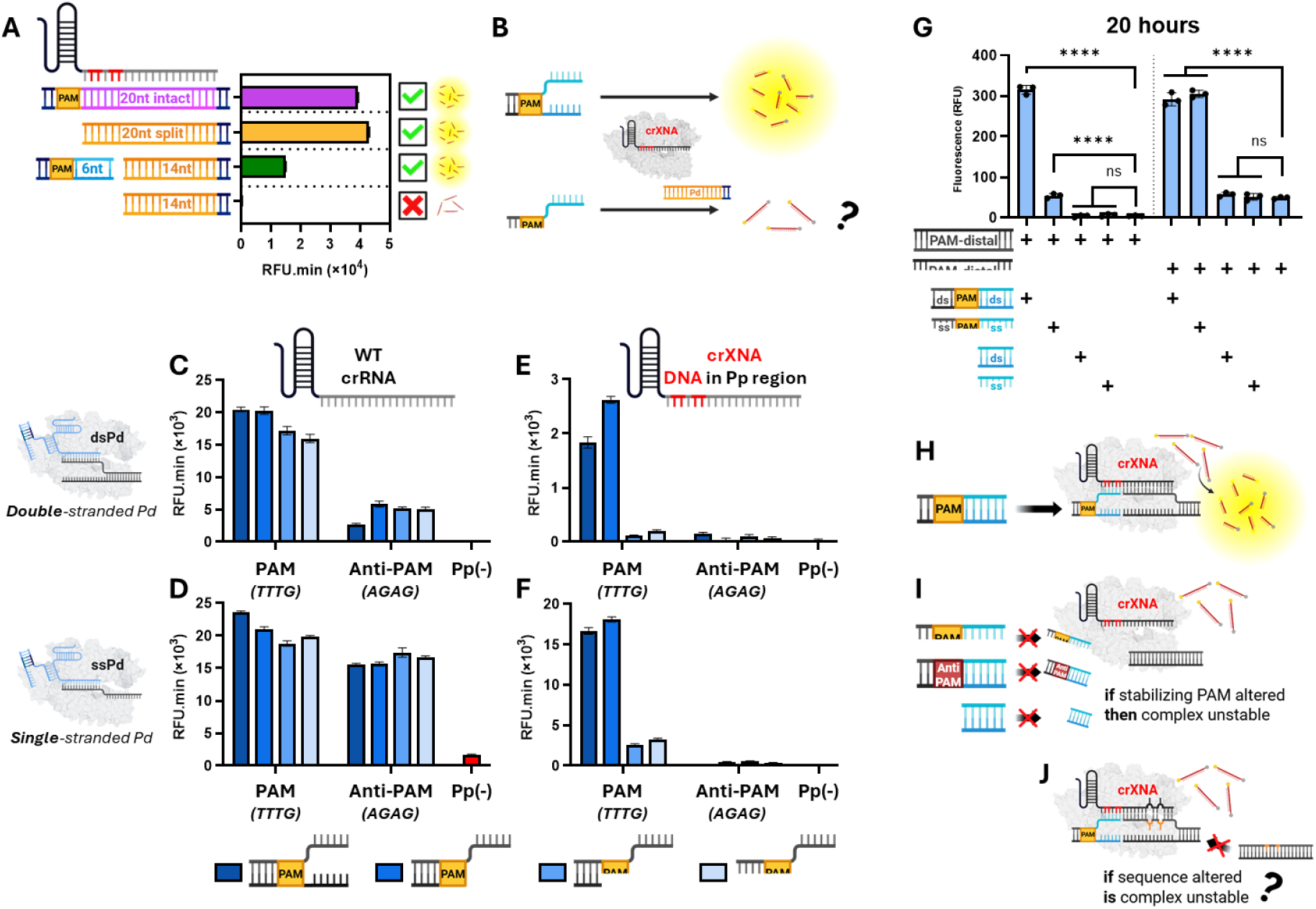
Improved substrate selectivity with chimeric guide. **(A)** Trans-cleavage activity of crXNA-guided Cas12a targeting a full-length protospacer with or without the PAM, and split activator with or without the Pp. Background-subtracted fluorescence intensity was measured over 2 hours at intervals of 200 seconds. Values represent total area under the curve (RFU.min) ± Standard error. **(B)** crXNA-mediated activation of Cas12a requires a double-stranded PAM in the context of a split activator. **(C-F)** crRNA-(left column) and crXNA-mediated (right column) activation of Cas12a was compared when activated with Pp of different structures and PAM sequences as well as double-stranded (top row) or single-stranded (bottom row) Pd activators. Background-subtracted fluorescence intensity was measured over 60 minutes, with readings every 220 seconds. Values represent total area under the curve (RFU.min) ± Standard error. **(G)** Trans-cleavage of the reporter was measured after a crXNA-guided Cas12a was provided a double-stranded Pd activator, various structures of the Pp activator, and incubated over 20 hours. Values represent mean background-subtracted fluorescence intensity ± SD. Analyses were performed using a one-way ANOVA with a multiple comparison test where ns = not significant with *p* > 0.05 and asterisks (\**p* ≤ 0.05, \*\**p* ≤ 0.01, \*\*\**p* ≤ 0.001, and \*\*\*\**p* ≤ 0.0001) denote significant differences. Graphical depictions of crXNA-guided Cas12a associating to a Pp and **(H)** activating when the PAM is properly structured and **(I)** not activating when the PAM is altered. This is likely because destabilizing hybridization in Pp-bound intermediate makes split activator complexes prone to collapse when other key interactions are altered. **(J)** If so, then introducing mismatches between the crXNA and the split activators may similarly destabilize the complexes. All experiments performed at n=3. Graphics created in https://BioRender.com

To explore the PAM’s relevance in maintaining the Pp-bound complexes (Figure S14B-E), we compared crRNA-(normal hybridization) and crXNA-mediated (altered hybridization) activation using various Pp structures and PAM sequences (PAM=TTTG vs. Anti-PAM=AGAG) (Figures 4C-F). We first targeted a double-stranded Pd whose non-target strand would need to be displaced by the Pp-bound Cas12a, making association more challenging. When guided by a crRNA, activity was slightly reduced when the PAM was single-stranded and significantly reduced under an “Anti-PAM” (Figure 4C, S15A, S15B), reflecting the importance of the PAM’s structure and sequence in associating the Pp and Cas12a (Figure 2D,2E). When targeting a more accessible, single-stranded Pd, crRNA-mediated activity was drastically enhanced (Figure 4D). Here, the structure and sequence of the PAM still affected the activation of Cas12a, though the overall activity was very strong (Supplementary Figures S15C, S15D). From this, we can infer that under conditions where the crRNA can hybridize more easily to its target (single-stranded Pd target), the PAM is less relevant. This is either because I) the PAM was involved in unwinding the Pd target, which is now unnecessary with a single-stranded Pd target, or II) the PAM is involved in stabilizing the Pp-bound state, making it very important when targeting harder-to-access double-stranded Pd targets, but comparatively less relevant when targeting single-stranded targets.

To test this, we used the crXNA to weaken the nucleic acid binding energy between the Pp and the Cas12a. We noted that the crXNA-mediated activity was strong when the PAM was double-stranded, and all but eliminated when the PAM was either single-stranded or contained the incorrect sequence (Anti-PAM=GAGA) (Figures 4E, S16A, S16B). When the Pd-target was single-stranded and therefore easier to hybridize, the crXNA-guided Cas12a was slightly active when using a single-stranded PAM, but did not tolerate an Anti-PAM (Figures 4F, S16C, S16D). The crXNA-guided Cas12a was very active on the single-stranded Pd target with the PAM, implying that the PAM is important for Pp-binding when hybridization is weak, but not very active when the PAM was poor. This supports the hypothesis that the role of the PAM is to stabilize the Pp-bound state, as opposed to improve unwinding of the Pd.

Cas12a’s activity can be modulated through selective truncations in the PAM’s duplex (Supplementary Figure S17). When guided by a crXNA, the final position of the PAM does not affect Cas12a’s activity (5’-TTT<u>G</u>-3’) (Supplementary Figure S17C, S17D). The removal of the adjacent thymine significantly reduces Cas12a activation, with no noticeable activity over baseline when two thymines are removed (Supplementary Figure S17E, S17F). Interestingly, the Lys595 residue in the PAM-interaction domain is inserted between these two positions in and is thought to be involved in stabilizing the PAM-binding channel.(32) Compromising this interaction significantly reduced Cas12a activity, supporting the hypothesis that the protein relies more heavily on PAM-bound interactions in the context of a split activator (Supplementary Figure S14D, S14E).

Having determined that the PAM’s structure could modulate the activity of a split activator system, we wondered how well this effect would persist over a long incubation. To test this, we incubated Cas12a with various DNA substrates over 20 hours. When targeting a double-stranded Pd, whose NTS would need to be displaced for the crXNA to hybridize, the double-stranded PAM yielded a strong signal (Figure 4G, S18A), while the single-stranded PAM was significantly less intense. When targeting a more easily accessible, single-stranded Pd, Cas12a activity was weaker in the presence of the single-stranded PAM (Supplementary Figure S18B), but still resulted in full cleavage after 20 hours (Figure 4G). When the PAM was absent, the presence or absence of the Pp did not affect crXNA-guided Cas12a activity (Figure 4G, S18A, S18D). In comparison, the crRNA which hybridizes normally to the Pp could associate to the PAMless Pp and activate Cas12a (Supplementary Figures S18C, S18D).

Taken together, we can see that destabilizing the hybridization between the Pp and the guide (crXNA) caused the Cas12a to rely more heavily on the PAM when associating to the Pp (Figure 4H). Alterations to the PAM were therefore poorly tolerated by Cas12a, making it unlikely for the protein to associate with both the Pp and Pd, significantly reducing activity (Figure 4I). In other words, weakening the stability of the complexes made them prone to collapse if any other key interactions were compromised, effectively making Cas12a selective for the structure of the PAM. This therefore led us to wonder if destabilizing the complexes through mismatches between the crXNA and the split activator would similarly make Cas12a specific for individual single-nucleotide polymorphisms (Figure 4J).

### Improved specificity with chimeric guide

When targeting an intact activator, Cas12a anneals its crRNA to the TS in a process known as R-loop elongation. This dynamic process is primarily driven by the electrostatic forces from DNA:RNA hybridization and DNA:DNA unpairing, with little intervention on the part of Cas12a.(7,42) As such, mismatches in the TS increase the dissociation kinetics of the growing R-loop, favoring its collapse and preventing Cas12a activation. However, this is not a perfect process and outside of its seed region (first ∼5 nt), Cas12a is not very specific to individual mismatches between its crRNA and the TS.(5)

To test if the less stable crXNA-guided Cas12a could be more specific to mismatches in the context of a split activator, we first investigated the effects of mutations along the Pp (Supplementary Figure S19). Individual mutations were installed along all six bases of the Pp and the first six bases of the control (intact) activator. As expected, the crRNA-guided Cas12a was not particularly specific to individual mutations in an intact activator (Supplementary Figure S19A). Interestingly, neither the crXNA targeting an intact activator (Supplementary Figure S19B), nor the crRNA targeting a split activator (Supplementary Figure S19C) improved the specificity of Cas12a. However, when a crXNA was used to target a split activator, all but the fifth position in the crRNA became more specific to individual mutations (Supplementary Figure S19D). This indicates that disfavoring stability of the Pp:crXNA duplex through base substitutions disproportionately affects the stability of the Pp:XNP complex, leading to improved specificity for individual mutations. However, the overall specificity was still relatively lackluster, possibly because the dsPAM in the Pp allowed the Pp:XNP to reassemble from the PAM-bound conformation after R-loop collapse, reducing the effects of mismatches in the duplex (Supplementary Figure S14D).

Since the Pd does not have a PAM, we reasoned that R-loop collapse along this fragment could result in improved specificity. Like before, we substituted individual positions in the Pd and an intact activator (positions 7-20) and used the new duplexes to activate Cas12a. When targeting an intact activator, the crRNA-guided Cas12a was not particularly specific (Figures 5A, S20). Weakening the hybridization along the seed region of the crXNA (Figures 5B, S21) or splitting the activator while using a crRNA (Figures 5C, S22) each noticeably improved the specificity of Cas12a in many positions, even after 90 minutes. In the former case, this may be because the seed-bound conformation (crXNA+intact) was less stable, favoring the dissociation of the Cas12a from the DNA duplex once the R-loop reverse propagated sufficiently far. In the case of the latter, the reverse propagation of the R-loop in the Pd likely resulted in the Pp:RNP completely dissociating from the Pd, as expected. Interestingly, combining both approaches all but eliminated activity in Cas12a (Figure 5D, S23). Based on these results, we hypothesize that the reduced stability at the Pp:crXNA interface promoted the reverse propagation of the R-loop in the Pd under non-complementary base-pairing. To determine if this improvement in specificity was effective in discriminating between transition and transversion mutations, we next prepared Pd duplexes with all possible combinations of mismatches (e.g. C:G, C:T, C:A, and C:C) for each of the four bases (e.g. A, C, G, and T) in the crRNA (Supplementary Figure S24A, S24B). Indeed, the crXNA-guided Cas12a demonstrated remarkable specificity against these substrates (Supplementary Figure S24C-J), with the interesting exception of a G:T mismatch at position 12 of the crRNA’s spacer (Supplementary Figure S24K, S24L). Furthermore, the activity of the crXNA-guided Cas12a on a split activator with two adjacent polymorphisms, is absent for all combinations of mutations, aside from the final 19/20 mutation (Supplementary Figure S25, S26).

**Figure 5:**
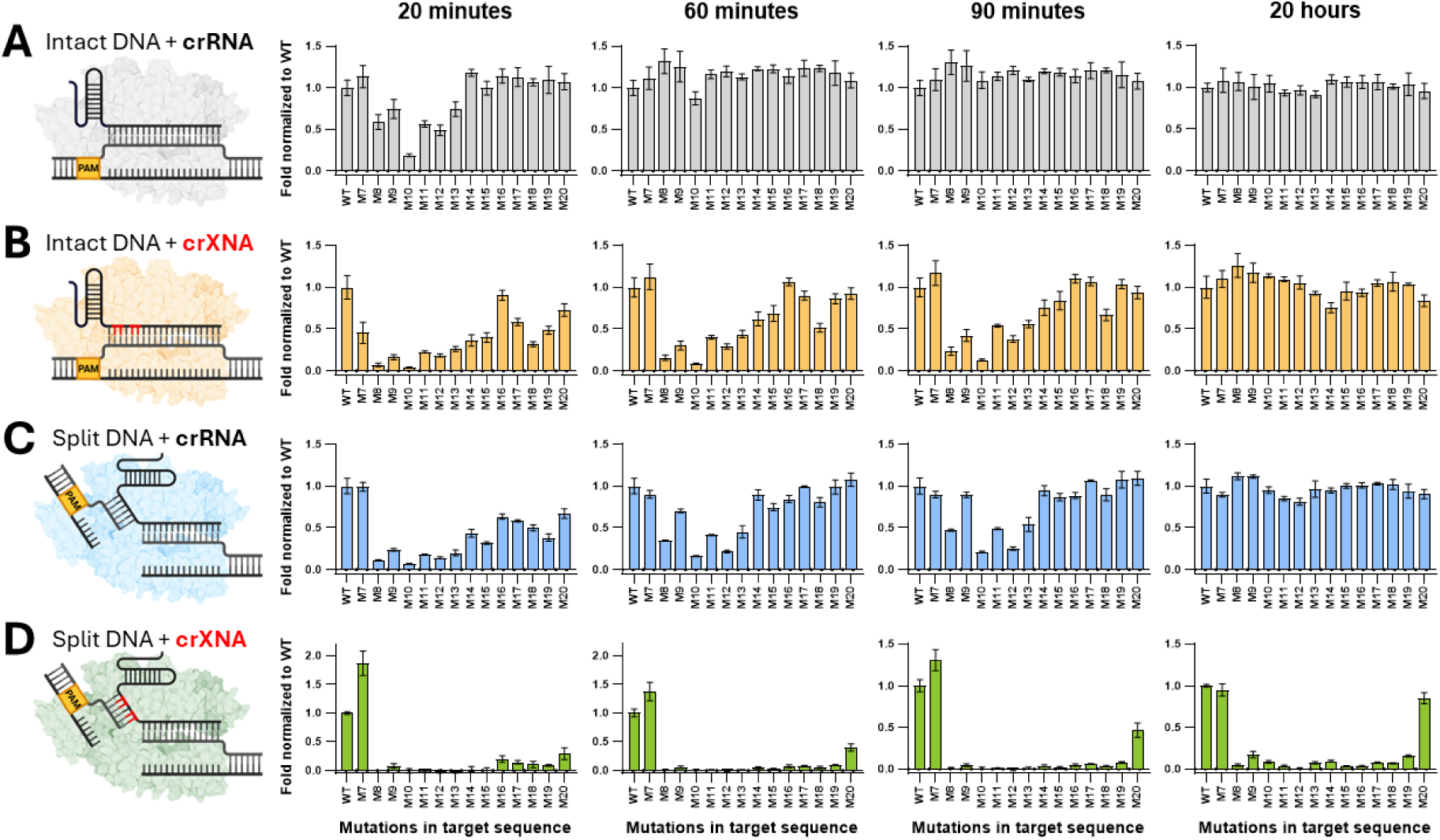
Activation of intact vs. split activator-based Cas12a reaction schemes on single-nucleotide polymorphisms (SNPs) in the Pd. An intact 20 nt protospacers was used to activate **(A)** a crRNA-guided (Grey) and a **(B)** crXNA-guided (Blue) Cas12a. This was compared to a split activator scheme with a Pp/Pd split at 6nt/14nt. When using split activators, the Cas12a was guided by either **(C)** a crRNA (Blue) or **(D)** a crXNA (Green). Values represent mean background-subtracted signal intensity normalized to complementary activator ± SD at different times. All experiments performed at n=3. Graphics created in https://BioRender.com

Finally, given that the crXNA-guided Cas12a was slow to activate on a split activator system, we sought to increase the speed at which the protein would activate. To do so, we tested a gradient of PEG-8000, a crowding agent. In so doing, we noticeably improved the activity of Cas12a over time (Supplementary Figure S27A), and crucially, Demonstrated that unlike with the crRNA, the crXNA maintained a very strong degree of specificity (Supplementary Figure S27B, S27C, S27D).

## DISCUSSION

In this study, we have explored the structural requirements of double-stranded split activator systems and how these components assemble towards the activation of CRISPR/Cas12a. First, we discovered that Cas12a readily activates even when its protospacer is separated from its PAM. Furthermore, this activity is maintained even when the protospacer lacks most but not all of the guide RNA’s seed region. This corroborates and builds upon a previous report which proposed that Cas12a can target split activators using frayed DNA extremities as surrogates for the PAM.(24) Indeed, we validated these findings and built on the understanding of this activity by demonstrating that the frayed DNA must complement some, but not all of the pre-structured seed in the crRNA.

Next, we thoroughly explored the assembly of the various components in the split activator system. We experimentally demonstrated that the role of the PAM is to mediate the assembly between the truncated PAM-proximal (Pp) and RNP, and that this mediation is based not only on the PAM’s sequence, but surprisingly also its structure. Given the strong evidence demonstrating the role of the PAM in mediating the co-localization between the Pp and the Cas12a, we propose that the PAM primarily favors the unwinding of the PAM-distal (Pd) by stabilizing the Pp:RNP complex, rather than by directly inducing structural rearrangements in the Cas12a. Contrary to previous literature, we additionally found that a Pd which does not induce notable trans-cleavage activity in the Cas12a can still associate to the protein.(24) We have therefore proposed a revised model detailing the association of the split activator system.

To further demonstrate the importance of the direct DNA:protein interactions in the context of a split activator, we selectively weakened the Pp:crRNA interface. By replacing the crRNA with a chimeric DNA/RNA guide, we forced the Cas12a to rely more heavily on its interactions with a properly duplexed PAM. This enhances our understanding of Cas12a by further showcasing the role of the PAM in immobilizing the Cas12a during target interrogation. Additionally, given the recent focus on single-stranded split activators, understanding the importance of the double-stranded PAM in associating the Cas12a and the TS can be leveraged to improve downstream applications such as logic gates and diagnostics, hence our work.

On top of increasing the stringency for the structural recognition of the Pp, destabilizing the Pp:RNP interaction significantly increased the burden on the complementarity between the Pd and the guide. This was leveraged to achieve near total specificity along all but the first and final bases in the Pd. This specificity remained extremely strong regardless of whether the system was used on transition or transversion mutations, with only one exception. This architecture has incredible potential for the field of diagnostics as detecting individual mutations in the Pd is inherently a PAM-less process, overcoming yet another of Cas12a’s major limitations.

In conclusion, our study provides a detailed breakdown for the activity profiles of a split double-stranded activator when using Cas12a with different truncations. The detailed analysis of how the complexes form will allow future work to improve the types of activity that can be leveraged using such an architecture, a feat we have proven possible by already making the specificity near perfect in PAM-less target sequences.

## ACKNOWLEDGEMENTS

Graphics for all figures were created with Biorender.com.

## AUTHOR CONTRIBUTIONS

**G.L.** Project administration, Conceptualization, Formal analysis, Methodology, Investigation, Validation, Visualization, Writing—original draft. **F.V.** Conceptualization, Formal analysis, Methodology, Investigation, Validation, Writing—original draft. **I.I.** Conceptualization, Supervision, Writing-review & editing. C.B.**, K.G., Y.L., J.R., A.C.** Writing-review & editing. **K.P.** Supervision, Visualization, Writing—review & editing. **J.P.T.** Resources, Supervision, Writing-review & editing.

## SUPPLEMENTARY DATA

Supplementary Figures S1-S27 and oligonucleotide sequences (Supplementary Table S1) are available at NAR online.

## DATA AVAILABILITY

All data supporting the findings of this study are available within the article and its online supplementary materials. Data supporting the plots are available upon reasonable request to the corresponding authors.

## REFERENCES

1. Barrangou, R., Fremaux, C., Deveau, H., Richards, M., Boyaval, P., Moineau, S., Romero, D.A. and Horvath, P. (2007) CRISPR provides acquired resistance against viruses in prokaryotes. Science, 315, 1709–1712.

2. Mojica, F.J., Diez-Villasenor, C., Garcia-Martinez, J. and Soria, E. (2005) Intervening sequences of regularly spaced prokaryotic repeats derive from foreign genetic elements. J Mol Evol, 60, 174–182.

3. Zetsche, B., Gootenberg, J.S., Abudayyeh, O.O., Slaymaker, I.M., Makarova, K.S., Essletzbichler, P., Volz, S.E., Joung, J., van der Oost, J., Regev, A., et al. (2015) Cpf1 is a single RNA-guided endonuclease of a class 2 CRISPR-Cas system. Cell, 163, 759–771.

4. Stella, S., Alcon, P. and Montoya, G. (2017) Structure of the Cpf1 endonuclease R-loop complex after target DNA cleavage. Nature, 546, 559–563.

5. Chen, J.S., Ma, E., Harrington, L.B., Da Costa, M., Tian, X., Palefsky, J.M. and Doudna, J.A. (2018) CRISPR-Cas12a target binding unleashes indiscriminate single-stranded DNase activity. Science, 360, 436–439.

6. Swarts, D.C., van der Oost, J. and Jinek, M. (2017) Structural Basis for Guide RNA Processing and Seed-Dependent DNA Targeting by CRISPR-Cas12a. Mol Cell, 66, 221–233 e224.

7. Strohkendl, I., Saifuddin, F.A., Rybarski, J.R., Finkelstein, I.J. and Russell, R. (2018) Kinetic Basis for DNA Target Specificity of CRISPR-Cas12a. Mol Cell, 71, 816–824 e813.

8. Swarts, D.C. and Jinek, M. (2019) Mechanistic Insights into the cis- and trans-Acting DNase Activities of Cas12a. Mol Cell, 73, 589–600 e584.

9. Teng, F., Li, J., Cui, T., Xu, K., Guo, L., Gao, Q., Feng, G., Chen, C., Han, D., Zhou, Q. et al. (2019) Enhanced mammalian genome editing by new Cas12a orthologs with optimized crRNA scaffolds. Genome Biol, 20, 15.

10. McCarty, N.S., Graham, A.E., Studena, L. and Ledesma-Amaro, R. (2020) Multiplexed CRISPR technologies for gene editing and transcriptional regulation. Nat Commun, 11, 1281.

11. Ling, X., Chang, L., Chen, H., Gao, X., Yin, J., Zuo, Y., Huang, Y., Zhang, B., Hu, J. and Liu, T. (2021) Improving the efficiency of CRISPR-Cas12a-based genome editing with site-specific covalent Cas12a-crRNA conjugates. Mol Cell, 81, 4747–4756 e4747.

12. Gier, R.A., Budinich, K.A., Evitt, N.H., Cao, Z., Freilich, E.S., Chen, Q., Qi, J., Lan, Y., Kohli, R.M. and Shi, J. (2020) High-performance CRISPR-Cas12a genome editing for combinatorial genetic screening. Nat Commun, 11, 3455.

13. Sanz Juste, S., Okamoto, E.M., Nguyen, C., Feng, X. and Lopez Del Amo, V. (2023) Next-generation CRISPR gene-drive systems using Cas12a nuclease. Nat Commun, 14, 6388.

14. Zhang, X., Wang, J., Cheng, Q., Zheng, X., Zhao, G. and Wang, J. (2017) Multiplex gene regulation by CRISPR-ddCpf1. Cell Discov, 3, 17018.

15. Wang, J., Lu, A., Bei, J., Zhao, G. and Wang, J. (2019) CRISPR/ddCas12a-based programmable and accurate gene regulation. Cell Discov, 5, 15.

16. Chen, S., Wang, R., Peng, S., Xie, S., Lei, C., Huang, Y. and Nie, Z. (2022) PAM-less conditional DNA substrates leverage trans-cleavage of CRISPR-Cas12a for versatile live-cell biosensing. Chem Sci, 13, 2011–2020.

17. Broughton, J.P., Deng, X., Yu, G., Fasching, C.L., Servellita, V., Singh, J., Miao, X., Streithorst, J.A., Granados, A., Sotomayor-Gonzalez, A. et al. (2020) CRISPR-Cas12-based detection of SARS-CoV-2. Nat Biotechnol, 38, 870–874.

18. Gootenberg, J.S., Abudayyeh, O.O., Lee, J.W., Essletzbichler, P., Dy, A.J., Joung, J., Verdine, V., Donghia, N., Daringer, N.M., Freije, C.A. et al. (2017) Nucleic acid detection with CRISPR-Cas13a/C2c2. Science, 356, 438–442.

19. Gootenberg, J.S., Abudayyeh, O.O., Kellner, M.J., Joung, J., Collins, J.J. and Zhang, F. (2018) Multiplexed and portable nucleic acid detection platform with Cas13, Cas12a, and Csm6. Science, 360, 439-444.

20. Rananaware, S.R., Vesco, E.K., Shoemaker, G.M., Anekar, S.S., Sandoval, L.S.W., Meister, K.S., Macaluso, N.C., Nguyen, L.T. and Jain, P.K. (2023) Programmable RNA detection with CRISPR-Cas12a. Nat Commun, 14, 5409.

21. Xi, X., Li, T., Huang, Y., Sun, J., Zhu, Y., Yang, Y. and Lu, Z.J. (2017) RNA Biomarkers: Frontier of Precision Medicine for Cancer. Noncoding RNA, 3.

22. Cruz, P.D., Wargowsky, R., Gonzalez-Almada, A., Sifontes, E.P., Shaykhinurov, E., Jaatinen, K., Jepson, T., Lafleur, J.E., Yamane, D., Perkins, J. et al. (2024) Blood RNA Biomarkers Identify Bacterial and Biofilm Coinfections in COVID-19 Intensive Care Patients. J Intensive Care Med, 39, 1071–1082.

23. Lamothe, G., Carbonneau, J., Joly Beauparlant, C., Vincent, T., Quessy, P., Guedon, A., Kobinger, G., Lemay, J.F., Boivin, G., Droit, A. et al. (2023) Rapid and Technically Simple Detection of SARS-CoV-2 Variants Using CRISPR Cas12 and Cas13. CRISPR J, 6, 369–385.

24. Huang, S., Lou, Y. and Zheng, L. (2024) Synergistic effect of split DNA activators of Cas12a with exon-unwinding and induced targeting effect. Nucleic Acids Res, 52, 11148–11157.

25. Yin, N., Yu, H., Zhang, L., Luo, F., Wang, W., Han, X., He, Y., Zhang, Y., Wu, Y., Pu, J. et al. (2025) Regulation of CRISPR trans-cleavage activity by an overhanging activator. Nucleic Acids Res, 53.

26. Iwe, I.A., Liu, F.X., Corsano, A., da Silva, S.J.R., Doucet, J., Singh, S., Lamothe, G., Zayani, R., Nguyen, J., Matthews, Q., et al. (2025) RAPID: Evaluation of Cas12a Protospacer Nicking and Chimeric Reporters for PAM-free RNA and DNA diagnostics. medRxiv, 2025.2007.2012.25331452.

27. Li, Q., Song, Z.L., Zhang, Y., Zhu, L., Yang, Q., Liu, X., Sun, X., Chen, X., Kong, R., Fan, G.C. et al. (2023) Synergistic Incorporation of Two ssDNA Activators Enhances the Trans-Cleavage of CRISPR/Cas12a. Anal Chem, 95, 8879–8888.

28. Hu, M., Cheng, X. and Wu, T. (2024) Modular CRISPR/Cas12a synergistic activation platform for detection and logic operations. Nucleic Acids Res, 52, 7384–7396.

29. Li, X., Wang, J., Cheng, X., Xu, Q., She, L. and Wu, T. (2025) A multi-functional synergistic platform of Cas12a split dsDNA activators. Chem Commun (Camb), 61, 6615–6618.

30. Lei, X., Ding, L., Yang, X., Xu, F., Wu, Y. and Yu, S. (2024) PAIT effect: Padlock activator inhibits the trans-cleavage activity of CRISPR/Cas12a. Biosens Bioelectron, 263, 116607.

31. He, W., Li, X., Li, X., Guo, M., Zhang, M., Hu, R., Li, M., Ding, S. and Yan, Y. (2024) Split activator of CRISPR/Cas12a for direct and sensitive detection of microRNA. Anal Chim Acta, 1303, 342477.

32. Yamano, T., Zetsche, B., Ishitani, R., Zhang, F., Nishimasu, H. and Nureki, O. (2017) Structural Basis for the Canonical and Non-canonical PAM Recognition by CRISPR-Cpf1. Mol Cell, 67, 633–645 e633.

33. Zgarbova, M., Otyepka, M., Sponer, J., Lankas, F. and Jurecka, P. (2014) Base Pair Fraying in Molecular Dynamics Simulations of DNA and RNA. J Chem Theory Comput, 10, 3177–3189.

34. Li, L., Pabit, S.A., Lamb, J.S., Park, H.Y. and Pollack, L. (2008) Closing the lid on DNA end-to-end stacking interactions. Appl Phys Lett, 92, 223901–2239013.

35. Maffeo, C., Luan, B. and Aksimentiev, A. (2012) End-to-end attraction of duplex DNA. Nucleic Acids Res, 40, 3812–3821.

36. Chen, J., Lin, X., Xiang, W., Chen, Y., Zhao, Y., Huang, L. and Liu, L. (2025) DNA target binding-induced pre-crRNA processing in type II and V CRISPR-Cas systems. Nucleic Acids Res, 53.

37. Grünweller, A. and Hartmann, R.K. (2007) Locked nucleic acid oligonucleotides: the next generation of antisense agents? BioDrugs, 21, 235–243.

38. Vester, B. and Wengel, J. (2004) LNA (locked nucleic acid): high-affinity targeting of complementary RNA and DNA. Biochemistry, 43, 13233–13241.

39. Kim, H., Lee, W.J., Oh, Y., Kang, S.H., Hur, J.K., Lee, H., Song, W., Lim, K.S., Park, Y.H., Song, B.S. et al. (2020) Enhancement of target specificity of CRISPR-Cas12a by using a chimeric DNA-RNA guide. Nucleic Acids Res, 48, 8601–8616.

40. Nakano, S., Kanzaki, T. and Sugimoto, N. (2004) Influences of ribonucleotide on a duplex conformation and its thermal stability: study with the chimeric RNA-DNA strands. J Am Chem Soc, 126, 1088–1095.

41. Yamano, T., Nishimasu, H., Zetsche, B., Hirano, H., Slaymaker, I.M., Li, Y., Fedorova, I., Nakane, T., Makarova, K.S., Koonin, E.V. et al. (2016) Crystal Structure of Cpf1 in Complex with Guide RNA and Target DNA. Cell, 165, 949–962.

42. Strohkendl, I., Saha, A., Moy, C., Nguyen, A.H., Ahsan, M., Russell, R., Palermo, G. and Taylor, D.W. (2024) Cas12a domain flexibility guides R-loop formation and forces RuvC resetting. Mol Cell, 84, 2717–2731 e2716.

